# eIF2α integrates proteotoxic signals both from ER and cytoplasm: Hsp70-Bag3 module regulates HRI-dependent phosphorylation of eIF2α

**DOI:** 10.1101/2021.04.13.439701

**Authors:** Shivani Patel, Santosh Kumar, Arkadi Hesin, Julia Yaglom, Michael Y. Sherman

**Affiliations:** Department of Molecular biology, Ariel University, Ariel, Israel

**Keywords:** Hsp70, Bag3, HRI, CHOP, eIF2α

## Abstract

The major heat shock protein Hsp70 has been implicated in many stages of cancer development. These effects are mediated by a scaffold protein Bag3 that binds to Hsp70 and links it to components of multiple cancer-related signaling pathways. Accordingly, the Hsp70-Bag3 complex has been targeted by small molecules, which showed strong anti-cancer effects. Here, our initial question was how JG-98, an allosteric inhibitor of Hsp70 that blocks its interaction with Bag3, causes cell death. Breast epithelial cells MCF10A transformed with a single oncogene Her2 showed higher sensitivity to JG-98 then parental MCF10A cells. RNA expression analysis showed that this enhanced sensitivity correlated with higher induction of the UPR genes. Indeed, depletion of the pro-apoptotic UPR responsive transcription factor CHOP significantly protected cells from JG-98. Surprisingly, only the eIF2α-associated branch of the UPR was activated by JG-98, suggesting that the response was not related to the ER proteotoxicity. Indeed, it was dependent on activation of a distinct cytoplasmic eIF2α kinase HRI. HRI-dependent phosphorylation of eIF2α was also activated by the cytoplasmic proteotoxicity via Hsp70-Bag3 complex, which directly associates with HRI. Dissociation of Hsp70-Bag3 complex led to Bag3-dependent degradation of HRI via autophagy. Therefore, eIF2α integrates proteotoxicity signals from both ER and cytoplasm, and the cytoplasmic response mediates cytotoxicity of the Hsp70-Bag3 inhibitors.

## Introduction

The major heat shock protein Hsp70 have been implicated in cancer development (1–3). Original hypothesis was that cancer cells experience permanent proteotoxicity stress due to aneuploidy, oxidative stress and other abnormalities, which dictates their stronger requirements for expression of molecular chaperones (non-oncogene addiction) compared to normal cells (4, 5). This hypothesis was generally refuted (6), which led to the idea of direct involvement of Hsp70 in signaling pathways that control major stages of cancer development, including initiation, progression and metastasis. Indeed, several publications demonstrated that knockout of Hsp70 leads to suppression of cancer development at various stages (depending on the cancer model) and these effects correlated with suppression of many signaling pathways involved in cancer development (7–11).

We and others previously reported that effects of Hsp70 on signaling pathways are mediated by the co-chaperone Bag3 that binds to the ATPase domain of Hsp70 and links it to components of various signaling pathways, like Src, Hif1, NF-kB, p53, Hippo and others (12–16). Knockout or depletion of Hsp70 or Bag3 affected activities of these pathways, which significantly influenced cancer cell physiology (13). With some of these pathways, direct interaction with Bag3 was reported, e.g. Src, Yes or Lats1 (13, 17, 18), while with other pathways, e.g. Hif1 or FoxM1, mechanisms of effects of Hsp70 and Bag3 remain unclear.

Very importantly, knockout of Hsp70 did not significantly affect normal organism development and physiology, while strongly suppressing proliferation and survival of cancer cells (19, 20). These and other findings made Hsp70 an attractive target for the anti-cancer drug design, and a number of attempts have been made to develop Hsp70 inhibitors as drug prototypes (21–29). Our collaborative work with the group of Jason Gestwicki reported a series of inhibitors of Hsp70, e.g. YM-01 or JG-98, that interact with an allosteric site at the ATPase domain of Hsp70 and freeze it in the ADP-bound form, which, in turn, leads to dissociation of Bag3 (13, 30). These inhibitors mimicked all effects of Hsp70 or Bag3 depletion on a series of signaling pathways both in cell culture and in tumor xenograft animal models (13, 30). Interestingly other Hsp70 inhibitors that do not cause dissociation of Bag3 did not show these effects on signaling pathways (13). Importantly, YM-01, JG-98 and more advanced inhibitors from this series, e.g. JG-231, demonstrated potent anti-cancer effects (10, 31–33). These effects were related to both direct suppression of growth of tumor cells and their death, as well as suppression of migration of tumor-associated macrophages into the tumor (31). Further investigation uncovered that effects of this class of Hsp70 inhibitors are entirely different from effects of Hsp90 inhibitors and are clearly related to dissociation of the Hsp70-Bag3 complex (10).

Because of the potential clinical importance of understanding anti-tumor effects of this class of Hsp70 inhibitors, here, we originally aimed to clarify how these compounds initiate cancer cell death. We observed enhanced sensitivity of transformed cells to JG-98 compared to untransformed cells, which correlated with stronger activation of multiple signaling pathways. This line of investigation led to understanding the central role of one of the branches of UPR in cell response to these Hsp70 inhibitors and uncovered a novel mechanism of cell signaling upon the buildup of abnormal polypeptides in cytoplasm.

## Results

Because of the high potential of targeting Hsp70-Bag3 complex for anti-cancer therapy, we sought to understand how disruption of the complex by JG-98 affects cell death. Previous analysis suggested that there are growth inhibition effects related to modulation of expression levels of proteins involved in the cell cycle, e.g. c-myc (10). Such inhibition of the cell cycle progression could be partially responsible for suppression of tumor growth by JG-98. On the other hand, there was clear cell death in part of cell population, and we decided to dissect the nature of this phenomenon. To assess effects of JG-98 on cell death, we compared untransformed MCF10A cells and MCF10A transformed with a single oncogene Her2. To transform, cells were infected with a retrovirus expressing neu mutant form of Her2 oncogene (Fig. S1), which significantly changed their phenotype from epithelial to more mesenchymal (not shown). In parallel, we compared gene expression data of control and transformed cell population and observed activation of the entire set of cancer-related pathways in the latter population (Fig. S2). Cell death was assessed by counting survived DAPI-stained cells, following incubation with 1uM JG-98 for 24 hours using Hermes imaging system (see Materials and Methods). Her2-dependent transformation of cells significantly enhanced their sensitivity to JG-98 (Fig. 1A), demonstrating cancer specificity of the response. The higher sensitivity of Her2-transformed cells to JG-98 was parallel to a stronger activation of stress and MAP kinase pathways by JG-98 (Fig. 1B), suggesting that overregulation of a signaling pathway may be responsible for the cell death effects. Interestingly, transformed cells demonstrated stronger responses not only to blocking Hsp70, but to a proteotoxic stress in general. Indeed, in transformed cells, we observed stronger activation of stress and MAP kinases following proteasome inhibition by MG132, compared to untransformed cells (Fig. 1C).

**Fig. 1.**
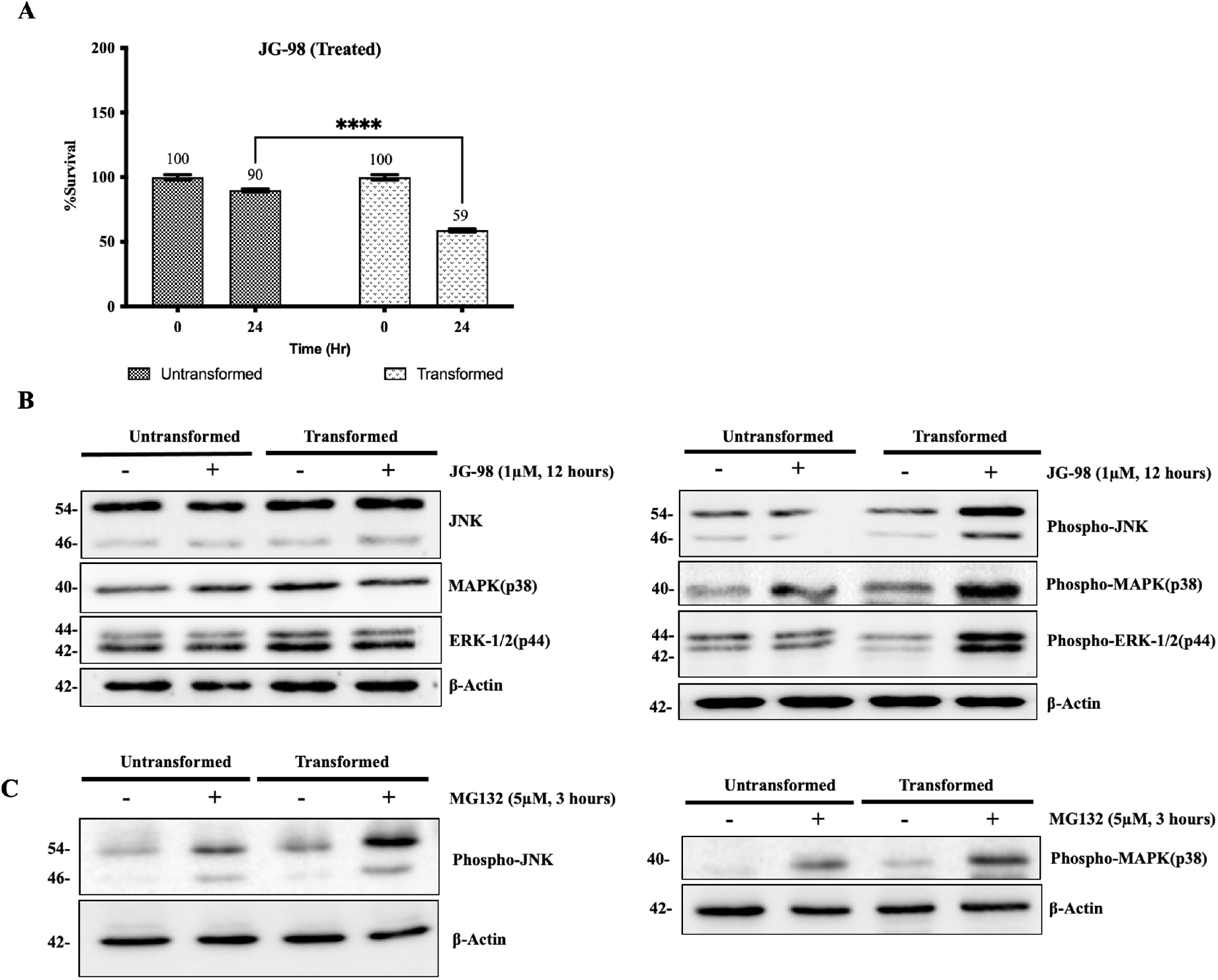
Her2 transformed cells showed enhanced sensitivity to JG-98 and stronger activation of stress kinases. **(A)** Transformation with Her2 oncogene enhances the sensitivity of the MCF10A cells to JG-98 treatment. MCF10A or Her2-transformed MCF10A cells were treated with 1μM JG-98 for 24 hours at 50% confluency or left untreated. Cell viability was quantified by counting of DAPI stained cells using Hermes Imaging systems. Respective control bar represents the relative percentage of cells at 24 hours after treatment compared to 100 percent at 0 hours of treatment. **(B)** MCF10A (Untransformed and Her2/Neu transformed) cells were treated with JG-98 at concentration of 1μM for 12 hours or left untreated. Levels of proteins were determined in cell lysates by immunoblotting with corresponding antibodies, JG-98 showed increase in the phosphorylation of phospho-JNK, phospho-MAPK(p-38) and phospho-ERK1/2(p44). Whereas, total JNK, MAPK(p-38), ERK-1/2 (p44) remain unchanged, see left panel. **(C)** MCF10A (Untransformed and Her2/Neu transformed) cells were treated with 5μM MG132 for 3 hours or left untreated, Levels of phospho-JNK and phospho-MAPK(p-38) were determined in cell lysates by immunoblotting with corresponding antibodies. Respective blots are representative of three biological replicates.

To obtain insights about the mechanism of the sensitivity, we assessed JG-98-induced activation of signaling pathways by the multiplexing immune-probing RPPA technology. RPPA provides information about expression levels and phosphorylation status of several hundred polypeptides components of major signaling pathways. Untransformed and Her2-transformed MCF10A cells were incubated either with 1uM of JG-98 or remained untreated and protein lysates were prepared and assessed by the RPPA analysis. A number of signaling proteins was up- or downregulated in response to JG-98 (Table S1), some of them belonging to defined pathways, suggesting a coordinate regulation. For example, we observed a strong increase in levels of mitochondrial chaperones Grp75, TRAP1 and TUFM, suggesting a mitochondrial stress response. RPPA analysis provided hints into additional pathways that could be regulated by Hsp70-Bag3 complex. For example, we observed downregulation of p90-RSK and alteration of the Akt-mTOR pathway culminating in downregulation of p70-S6K and corresponding reduction of phosphorylation of S6 ribosomal protein. The latter finding is consistent with previous reports of downregulation of Akt in response to depletion of Hsp70 or Bag3. We also observed upregulation of a series of proteins related to energy metabolism and transcription regulators (Table S1). Among downregulated proteins, we found components of the cell cycle belonging to transition through the G1 phase, including p-RB and cyclin D3. Downregulation of these genes could be responsible for the cell cycle effects of JG-98. However, in relation to cell death, the picture remained unclear. We observed downregulation of the pro-apoptotic Puma, and on the other hand of the anti-apoptotic pS474-Akt2.

Since RPPA did not provide clear insights into the JG-98-induced toxicity, we performed RNAseq analysis. Untransformed and Her2-transformed MCF10A cells were incubated either with 1uM JG-98 for 12 hours or remained untreated and RNA was isolated and sent for sequencing. After normalization of the results, we performed GSEA pathway analysis (Table 1). As expected, we found that pathways related to the cell cycle changes, e.g. E2F targets, G2/M and myc targets, were significantly downregulated. Among significantly upregulated pathways, we found UPR (Fig. 2A), which was represented by a set of genes, including ATF3, ATF4, CHAC1, CHOP (DDIT3) and others (Fig. 2B). Because of the known role of CHOP in apoptosis(41–47), we suggested that activation of this pathway may play an important role in induction of cell death by JG-98. Interestingly, expression of most of the UPR genes was stronger upregulated by JG-98 in Her2-transformed cells compared to untransformed MCF10A (Fig. 2B), which further supported the hypothesis of the role of UPR in JG-98-induced cell death, since the transformed cells were more sensitive to the drug. To test if CHOP is involved in the cell death caused by JG-98, we compared effects of CHOP depletion by siRNA on the MCF10A cell death in response to the drug. Control and CHOP-depleted cells were exposed to 1uM of JG-98 for 24 hours, and cells death was assessed by cell counting using an automated imaging system Hermes. Indeed, depletion of CHOP significantly suppresses the JG-98-induced cell death (Fig. 2C,D).

**Table 1.** Pathways up- and downregulated by JG-98 in untransformed and transformed MCF10A cells. FDR values are shown for each pathway.

| **Pathways upregulated** | **Untransformed**<br>**FDR-q.value** | **Transformed**<br>**FDR-q.value** |
| --- | --- | --- |
| HALLMARK_TNFA_SIGNALING_VIA_NFKB | 0.005 | 0.00 |
| HALLMARK_IL6_JAK_STAT3_SIGNALING | 0.023 | 0.095 |
| HALLMARK_UNFOLDED_PROTEIN_RESPONSE | 0.035 | 0.006 |
| HALLMARK_INFLAMMATORY_RESPONSE | 0.143 | 0.062 |
| HALLMARK_P53_PATHWAY | 0.396 | 0.077 |
| HALLMARK_APOPTOSIS | 0.349 | 0.097 |
| HALLMARK_HEME_METABOLISM | 0.378 | 0.143 |
| HALLMARK_PEROXISOME | 0.617 | - |
| HALLMARK_HEDGEHOG_SIGNALING | 0.578 | 0.322 |
| HALLMARK_KRAS_SIGNALING_DN | 0.546 | 0.778 |
| HALLMARK_KRAS_SIGNALING_UP | 0.735 | 0.069 |
| HALLMARK_TGF_BETA_SIGNALING | 0.999 | 0.658 |
| **Pathways Downregulated** | **Untransformed**<br>**FDR-q.value** | **Transformed**<br>**FDR-q.value** |
| HALLMARK_E2F_TARGETS | 0.127 | 0.001 |
| HALLMARK_G2M_CHECKPOINT | 0.150 | 0.015 |
| HALLMARK_MYC_TARGETS_V2 | 0.157 | 0.010 |
| HALLMARK_MYC_TARGETS_V1 | 1.000 | 0.133 |
| HALLMARK_TGF_BETA_SIGNALING | 1.000 | - |

**Fig. 2.**
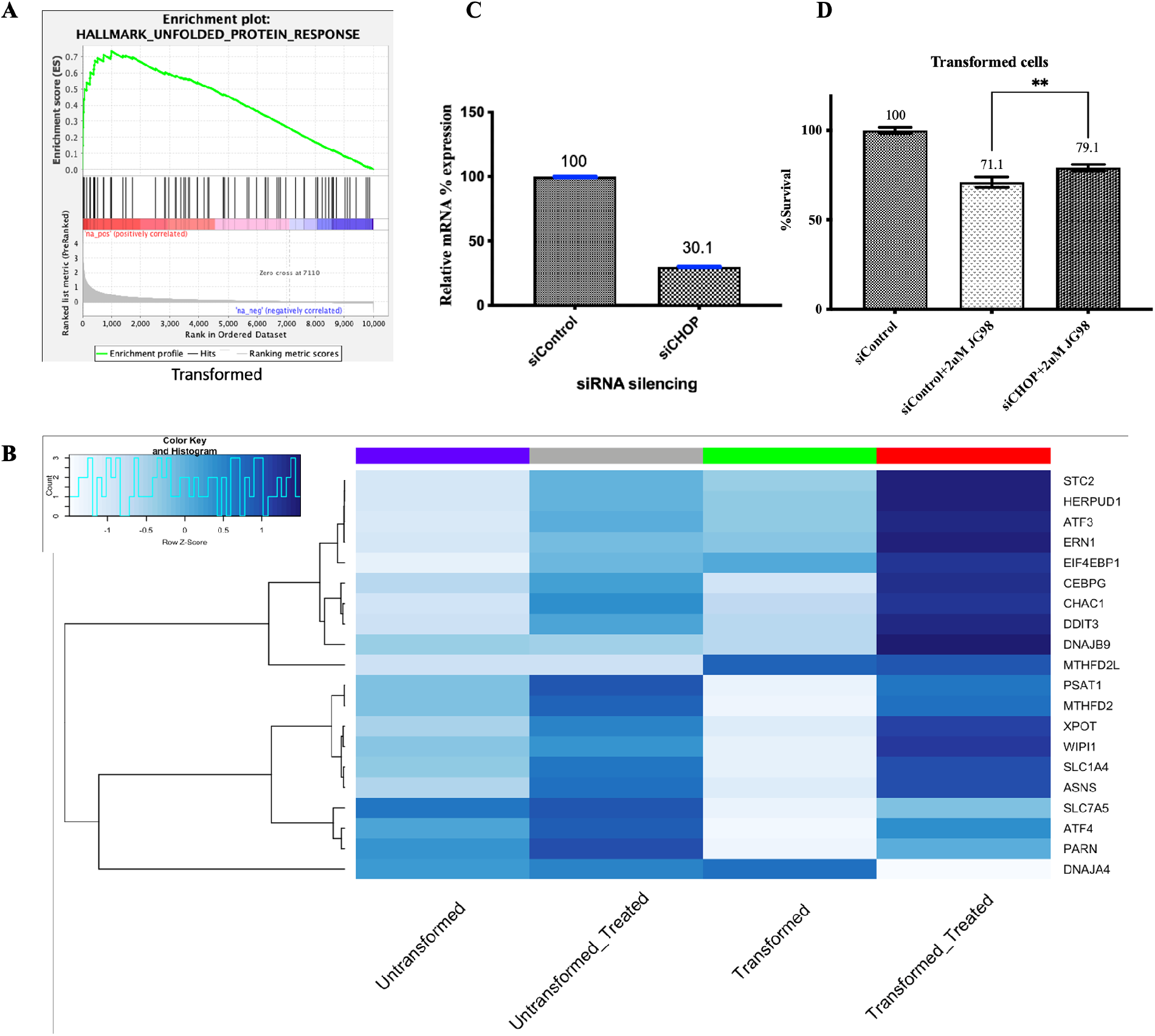
JG-98 triggers ATF4-CHOP axis of the unfolded protein response. Transcriptome analysis was performed with untransformed and transformed MCF10A cells treated with 1μM JG-98 treatment for 12 hours or left untreated. **(A)** UPR shows significant upregulation following JG-98 treatment using GSEA analysis of RNAseq. **(B)** Hallmark genes of unfolded protein response are significantly enriched in transformed cells in response to JG-98 treatment compared to untransformed cells. Color key represents the raw z-score. **(C)** CHOP (DDIT3) was depleted using siRNA and level of mRNA was quantified in comparison to siControl through quantitative PCR. **(D)** CHOP depletion reduces cell death in response to JG-98. NeuT (Her2/Neu Transformed MCF10A cells) were transfected with siControl and siCHOP to silence CHOP followed by 2μM JG-98 treatment for 24 hours or left untreated. Cell survival was evaluated by Hermes Imaging of DAPI-stained cells. Statistical analysis was performed by one-way ANOVA using Graphpad (v9).

Upregulation of the set of UPR genes suggested that in response to JG-98, the upstream regulator of the pathway a translation initiation factor eIF2α could become phosphorylated, and thus inactivated, which we indeed observed (Fig. 3A). As with other signaling pathways, phosphorylation of eIF2α was stronger activated by JG-98 in Her2-transformed cells (Fig. 3A). Activation of UPR suggests that JG-98 could cause a build-up of abnormal proteins in the endoplasmic reticulum. This effect would be consistent with possible inhibition of the ER resident Hsp70 family member BiP, which could be a direct target of JG-98. However, this possibility appears not to be the case. eIF2α phosphorylation represents only one out of three branches of the UPR response, which is regulated by the ER-associated PERK1 kinase (48). All three branches of UPR respond to titration of BiP by damaged polypeptides (49). Two other branches are controlled by other ER-associated proteins IRE1 and ATF6 (50). These two branches regulate expression of distinct sets of genes, e.g. ER chaperones. Surprisingly, among genes upregulated by JG-98, there were no ER chaperones or other genes regulated by either IRE1 or ATF6. Accordingly, it was unlikely that effects of JG-98 could be associated with inhibition of BiP and accumulation of the abnormal proteins in ER. Therefore, phosphorylation of eIF2α was probably triggered not by PERK1 (which is coregulated with IRE1 and ATF6) but by a distinct kinase. There are four known kinases that can phosphorylate eIF2α, including PERK1, a heme-responsive kinase HRI, an amino acid starvation regulated kinase GCN2, and a double strand RNA kinase PKR (51, 52). An important insight to distinguish between these kinase effects came from the GSEA analysis that showed a significant upregulation of the heme metabolism pathway in response to JG-98 (Fig. 3B), suggesting that HRI is activated.

**Fig. 3.**
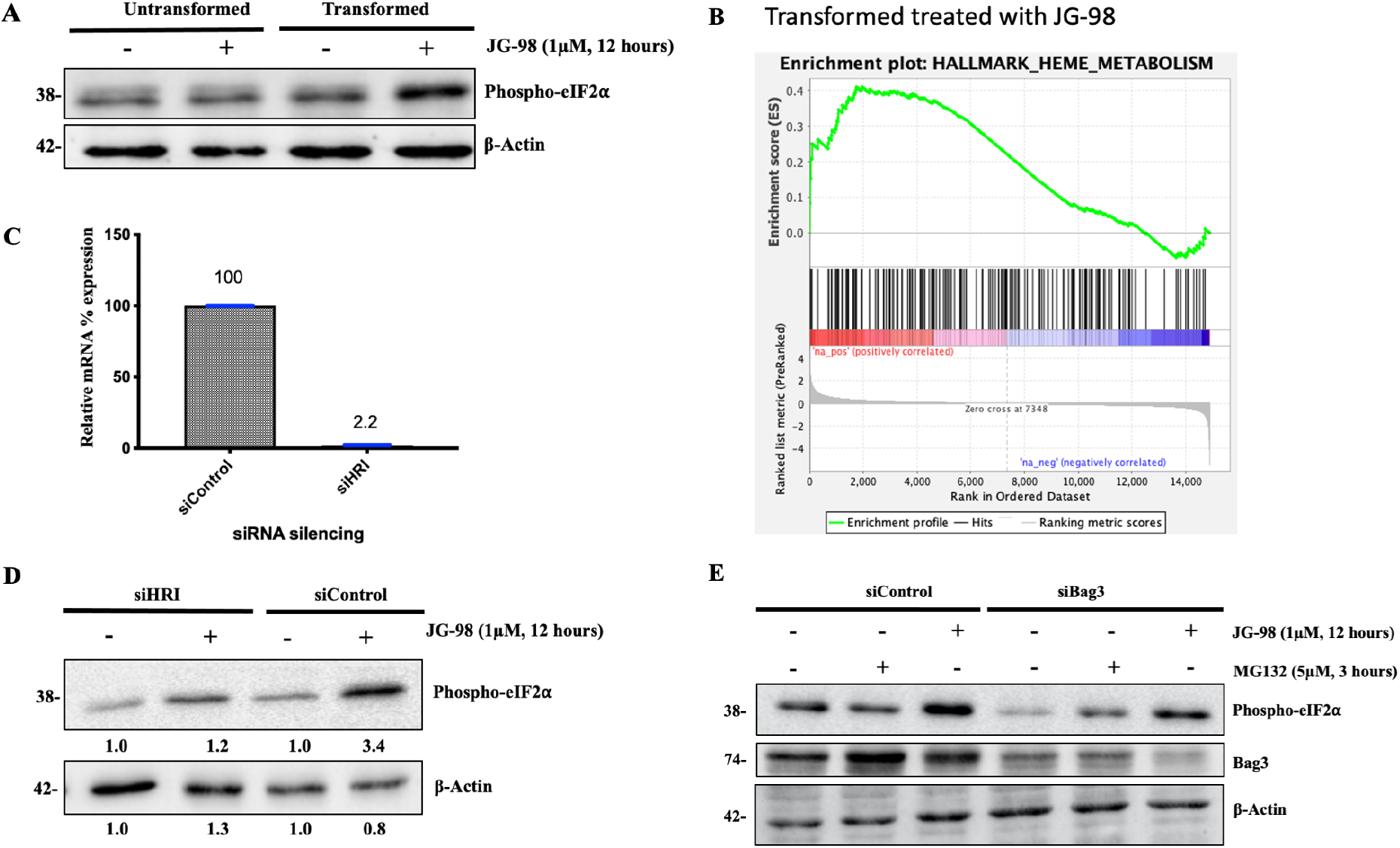
Phosphorylation of translation initiation factor eIF2α is mediated by Hsp70-Bag3-HRI complex in response to JG-98. **(A)** Treatment by JG-98 stronger stimulates phosphorylation of eIF2α in Her2/Neu transformed cells compared to untransformed cells. Cells were treated with 1μM JG-98 for 12 hours or left untreated. Levels of phospho-eIF2α were determined in cell lysates by immunoblotting with corresponding antibody. **(B)** GSEA analysis showing upregulation of the Heme metabolism pathway in response to JG-98 treatment of transformed cells. **(C)** Depletion of HRI was carried out through respective siRNA for 48 hours, as quantified through RT-PCR. **(D)** Depletion of HRI led to significant suppression of eIF2α phosphorylation in presence of MG132. **(E)** Depletion of Bag3 significantly reduced phosphorylation of eIF2α in the presence of JG-98 and MG132. Cells were transfected with siBag3 or siControl and further were treated by JG-98 (1uM, 12 hours) or MG132 (5uM, 3hours) or left untreated. Levels of phospho-eIF2α were determined in cell lysates by immunoblotting with corresponding antibody and normalized with beta-actin for quantification. Results are representative of three biological replicates.

To test for the role of HRI in phosphorylation of eIF2α following JG-98 treatment, we knocked down HRI using corresponding siRNA (Fig. 3C) and tested for the ability of JG-98 to facilitate phosphorylation of eIF2α. Indeed, depletion of HRI led to a significant downregulation of eIF2α phosphorylation in response to JG-98 (Fig. 3D). Importantly, depletion of a distinct eIF2α kinase GCN2 (Fig. S3) did not affect phosphorylation of eIF2α following treatment with JG-98 (Fig. S4). Therefore, HRI appears to be the major kinase that controls phosphorylation of eIF2α upon targeting Hsp70 by JG-98. These data suggest that HRI is controlled by the Hsp70-Bag3 module. Indeed, depletion of Bag3 using the corresponding siRNA also reduced phosphorylation of eIF2α under these conditions (Fig. 3E).

Since the major role of the Hsp70-Bag3 module in signaling is transmitting signals about the buildup of abnormal proteins in the cytoplasm to a variety of pathways, we suggested that this module transmits the proteotoxicity signals to HRI-eIF2α axis. Accordingly, we tested if triggering proteotoxicity by the proteasome inhibition can activate this axis in a Bag3-dependent manner. MCF10A cells were exposed to the proteasome inhibitor MG132 and eIF2α phosphorylation assessed either in control cells or cells depleted of HRI or Bag3. In fact, we observed that depletion of either HRI or Bag3 significantly suppressed phosphorylation of eIF2α in response to proteasome inhibition (Fig. 3E and 4A). Therefore, the proteotoxic stress in the cytoplasm can signal to phosphorylate eIF2α via Hsp70-Bag3-HRI axis. This conclusion was in line with previous report that HRI can associate with Hsp70 (53, 54).

The role of Hsp70-Bag3 in regulation of HRI suggested that the latter can associate with Bag3. To test this possibility, we expressed in cells 6His-tagged Bag3, and pulled it down together with the associated proteins either from naïve cells or cells treated with MG132. HRI was clearly pulled down in this experiment, and addition of MG132 did not significantly affect its association with Bag3 (Fig. 4B), indicating that proteotoxic stress regulates HRI in a way that does not involve its association/dissociation from Bag3. Interestingly, we could not see eIF2α in these pulldowns (not shown), suggesting that association of HRI and Bag3 with it is transient.

**Fig. 4.**
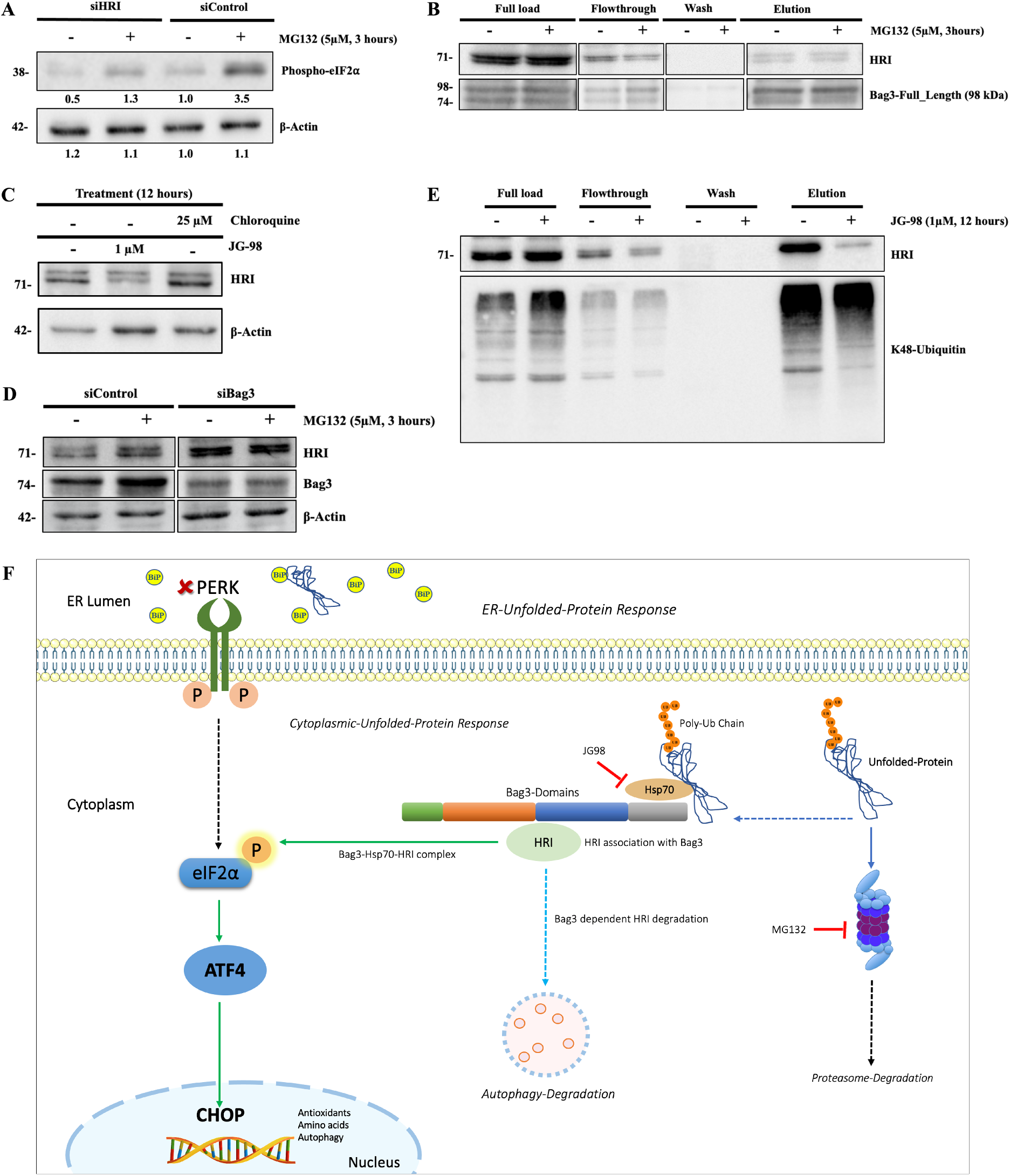
Proteotoxic stress in the cytoplasm signal to phosphorylate eIF2α via Hsp70-Bag3-HRI axis. **(A)** Depletion of HRI reduces phosphorylation of eIF2α in response to JG-98. **(B)** HRI associates with Bag3 in the pulldown assay. Association of HRI with Bag3 was assessed by expressing 6x-His tagged Bag3-Full length and pulled it down together with associated proteins (HRI) from naïve cells and cells treated with MG132 for 12 hours, **(C)** HRI levels are increased in the presence of the autophagy inhibitor hydroxychloroquine. **(D)** HRI levels are increased upon Bag3 depletion. **(E)** Hsp70-Bag3 complex link ubiquitinated proteins with HRI. Disruption of Hsp70-Bag3 complex with JG-98 causes dissociation of HRI from polyubiquitinated proteins. Ubiquitinated proteins were pulled down (see Materials and Methods) from cells treated or untreated with JG-98 and HRI levels in the pulldowns were measured by immunoblotting. **(F)** Schematic diagram demonstrates that eIF2α integrates proteotoxic signals both from ER and cytoplasm, the cytoplasmic response is mediated by HRI-Bag3. An additional Bag3 dependent pathway is HRI degradation via autophagy.

Direct association of HRI with Bag3 was in line with our observation that HRI can be degraded by autophagy. Indeed, incubation with the autophagic inhibitor hydroxychloroquine for 12 hours significantly increased the level of HRI (Fig. 4C), while similar incubation with the proteasome inhibitor MG132 did not affect the level of HRI (Fig. 4D). The autophagic degradation likely was Bag3-dependent, since Bag3 depletion led to similar increase in the HRI level (Fig. 4D). Therefore, it seems that Bag3 mediates both HRI-dependent phosphorylation of eIF2α and HRI degradation via the autophagic pathway.

Overall these data suggested that regulation of HRI activity by the buildup of abnormal proteins in the cytosol is mediated by the Hsp70-Bag3 module. Accordingly, Hsp70-Bag3 should link HRI with the abnormal ubiquitinated proteins. To test this model, we pulled down ubiquitinated proteins from the cells using an affinity purification with the UBA domain of ubiquilin-1 and immunoblotted with anti-HRI antibody. Indeed, a significant fraction of HRI associated with polyubiquitinated proteins (Fig. 4E). Importantly, addition of JG-98, which disrupts Hsp70-Bag3 complex dramatically reduced association of HRI with polyubiquitinated proteins (Fig. 4E), indicating that Hsp70-Bag3 mediates this interaction. Therefore, while BiP-Perk1 interaction mediates phosphorylation of eIF2α upon accumulation of abnormal proteins in ER, Hsp70-Bag3-HRI interaction mediates phosphorylation of eIF2α upon accumulation of abnormal proteins in cytoplasm.

## Discussion

Here, we first addressed how inhibitors that break Hsp70 interaction with Bag3 affect cancer cell death. We observed higher sensitivity to JG-98 of mammary epithelial cells transformed with a single oncogene Her2 than parental epithelial cells. This enhanced sensitivity was associated with stronger activation of stress and MAP kinases, suggesting that some of the pathways may be involved in the JG-98-induced cell death. Indeed, RNAseq followed by the pathway analyses predicted activation of UPR-related genes triggered by phosphorylation and inhibition of the translation initiation factor eIF2α. Though this activation was seen even in normal cells treated with JG-98, in transformed cells this response was significantly stronger. Since one of the genes in this pathway is a proapoptotic transcription factor CHOP, we suggested that its activation could contribute to the JG-98-induced cell death. Indeed, depletion of CHOP significantly protected cells from JG-98 treatment.

We were puzzled by the lack of activation of other branches of UPR by JG-98, strongly suggesting the lack of ER proteotoxicity under these conditions. Accordingly, we suggested that there might be a distinct pathway activated by JG-98 that leads to phosphorylation of eIF2α. Indeed, a cytosolic kinase HRI significantly contributed to eIF2α phosphorylation under these conditions. HRI was associated with Bag3, and its level was regulated by Bag3-dependent autophagy.

Prior works reported that HRI is responsible for phosphorylation of eIF2α in response to proteasome inhibitors (40). Here, we confirm this observation, and demonstrate that this response to proteotoxicity in cytosol is mediated by Bag3, since Bag3 depletion significantly reduced the levels of eIF2α phosphorylation. In this pathway, Hsp70-Bag3 linked HRI with abnormal ubiquitinated polypeptides, since disruption of the Hsp70-Bag3 complex by JG-98 led to dissociation of HRI from the ubiquitinated species.

Overall, this work uncovered a novel Hsp70-Bag3-HRI pathway that detect the buildup of abnormal proteins in the cytosol and regulates phosphorylation of eIF2α. In turn, eIF2α integrates proteotoxicity signals both from ER and cytosol (Fig. 4F).

### Experimental procedures

#### Reagents and Antibodies

JG98 was purchased from AdooQ BIOSCIENCE, MG132 from Biomol, 6% formaldehyde and DAPI from Sigma, Blasticidin from Thermo Fisher Scientific. Antibodies against, SAPK/JNK (9252S), phospho-JNK (4668S), p38-MAPK(8690S), phospho-p38-MAPK (4511S), p44/42-MAPK (9102S), Phospho-p44/42-MAPK(4370S), Phospho-eIF2α(Ser51)-(3597S) were from Cell Signaling Technology; EIF2AK1 (HRI)-ab28530, was from abcam, anti-Bag3 (10599-1-AP, Proteintech) and β-actin (NB600-501, Novus biologicals), α-K48 Ubiquitin linkage antibody (A101-050), Recombinant Human His6-Ubiquilin-1 Tandem UBA (UBE110) were from Boston Biochem.

#### Constructs and Oligonucleotides

The retroviral expression vector pCXbsr containing Her2/NeuT was described previously (34). For pull-down experiments we used pcDNA3.1-based plasmids used for overexpression of N-terminally His-tagged Bag3 Full length described previously (35, 36).

We used the following siGENOME siRNAs purchased from Dharmacon (Supplement table): nontargeting siRNA no. 5, BAG3 (D-011957-01), GCN2 (D-016634-02), DDIT3 (M-017609-01), HRI (M-006968-00).

#### Real Time PCR analysis

Total RNA was isolated from cells using Qiagen RNeasy Plus Mini kit (Cat. #74134;GmbH). The qScript cDNA Synthesis Kit (Quanta Bio, Cat. #95047-100) was used to convert 1ug mRNA into cDNA. qRT–PCR was performed using the PerfeCTa SYBR Green FastMix ROX (Quanta Bio, Cat. #95073-012) according to the manufacture’s protocols; expression levels of -Actin were used as an internal control. Real-time analysis was performed with a AriaMx Real-Time PCR System (Agilent Technologies) instrument using the primers (Supplement table) from Integrated DNA technologies (IDT), Belgium.

#### Cell culture

MCF10A (human breast epithelial) cells were grown in DMEM/F-12 1:1 medium supplemented with 5% horse serum (Cat. #04-0041A, BI-Biologicals), 20 ng/mL epidermal growth factor (Cat. #AF-100-15,Peprotech, 1 mg), 0.5 μg/mL hydrocortisone(Cat. #H-0888, Sigma), 10 μg/mL human insulin (Cat. #I9278, Sigma), and 100 ng/mL cholera toxin (Cat. #C-8052, Sigma). HEK293T cells were grown in DMEM high glucose with 10% Fetal Bovine Serum (FBS, Cat. #04-007-1A, BI-Biologicals). All cultures were supplemented with L-glutamine (Cat. #03-020-1B, BI-Biologicals), L-alanyl-L-glutamine (Cat. #03-022-1B, BI-Biologicals), penstrep (Cat. #03-031-1B, BI-Biologicals), and were grown in a humidified incubator at 37°C and 5% CO2 (16). Cell cultures were checked for mycoplasma contamination routinely at six-week intervals.

#### Virus Preparation and infection

Retroviral expression vector pCXbsr expressing HER2/Neu gene under blasticidin (Cat. #A3784,0005, PanReac AppliChem) was used. Only pCXbsr without any insert was used as an ‘empty or E’ vector control. Retroviruses were produced as reported before (34, 37). Briefly, HEK293T cells were co-transfected with plasmids expressing retroviral proteins Gag-Pol, Vesicular somatitis virus G glycoprotein and our gene of interest or enhanced green fluorescent protein (37) using Lipofectamine 3000 (Cat. #L3000015,Invitrogen, Boston, USA). After 48 hours of transfection, supernatants containing the retroviral particles were collected, aliquoted, and frozen at -80° C until use. Cells were infected with diluted supernatant in the presence of 8μg/mL Polybrene overnight, and were selected with blasticidin (10μg/mL) 48 h after infection. Retroviral vectors expressing enhanced green fluorescent protein was used as infection efficiency indicator: virus dilution was used to achieve approximately 90% of infected cells being GFP positive 2 days after infection. Transformation of Neu/Her2 was also quantified through qPCR represented by Cq(ΔR) values (Fig. S1).

#### Primers and siRNA

M-Her2_fw: 5’-AACTGCAGTCAGTTCCTCCG-3’ (Ref# 223747403)

M-Her2_rev: 5’-GTGCTTGCCCCTCACATACT-3’ (Ref# 223747404)

huHRI_For: 5’-ACCCCGAATATGACGAATCTGA-3’ (Ref# 226359069)

huHRI_Rev: 5’-CAAGTGCTCCAGCAAAGAAAC-3’ (Ref# 226359070)

DDIT3_F: 5’-GAACGGCTCAAGCAGGAAATC-3’ (Ref# 226024031)

DDIT3_R: 5’-TTCACCATTCGGTCAATCAGAG-3’ (Ref# 226024032)

HuGCN2-F: 5’-TGGTAAACATCGGGCAAACTC-3’ (Ref# 226265602)

HuGCN2-R: 5’-GGACCCACTCATACAACAAGA-3’ (Ref# 22626503)

ACTB Human-For: 5’-CACCATTGGCAATGAGCGGTTC-3’ (Ref# 225252514)

ACTB Human-Rev: 5’-AGGTCTTTGCGGATGTCCACGT-3’ (Ref# 225252513)

siRNA sequences:

siBag3: GCAAAGAGGUGGAUUCUAA (D-011957-01-0005)

siCHOP-1: AAGAACCAGCAGAGGUCACUU (CTM-595156, Oligo ID: ALEIA-000013)

Human HRI: CUGAUUAAGGGUGCAACUAUU (CTM-595153, Oligo ID: ALEIA-000007)

Human GCN2: CACCGUCAAGAUUACGGACUU (CTM-595154, Oligo ID: ALEIA-000009)

siControl (Scrambled): AGGUAGUGUAAUCGCCUUU (CTM-595155, Oligo ID: ALEIA-595155)

#### Immunofluorescence

Cells were fixed at room temperature (RT) using 4% paraformaldehyde for 5 min and permeabilized at RT in 0.2% Triton X-100 in PBS for 10 min. Cells were washed two times with PBST (1xPBS,0.05%Tween) and incubated with DAPI (1:10000) for 5 min at RT, followed by three additional washes with PBST. Imaging was performed using Hermes Wiscon Imaging System (IDEA Bio-Medical Ltd.) and image analysis was performed using inbuilt software package system (Athena Wisoft, Ver: V1.0.10).

#### Immunoprecipitation and Immunoblotting

Cells were lysed with lysis buffer (50mM Tris-HCl (pH 7.4), 150mM NaCl, 1% Triton X-100, 5mM EDTA, 1mM Na3VO4, 50mM β-glycerophosphate, 50mM NaF) supplemented with Protease Inhibitor Cocktail (Cat. #P8340, Sigma) and Phenylmethylsulfonyl fluoride (PMSF). Samples were adjusted to have equal concentration of total protein and subjected to PAGE electrophoresis followed by immunoblotting with respective antibodies (16).

#### Pull-down experiments

For pull-down analysis of Bag3 and associated proteins, HEk293T cells were transfected in 10cm dishes with full length Bag3 construct, treated with different conditions. The cells were washed with DPBS, fixed with 1.2% formaldehyde for 10 minutes at room temperature, then Tris-HCl (pH 7.4) was added to 50mM final concentration which was followed by a wash with 50mM Tris-HCl (pH 7.4) in DPBS at 4°C. The cells were lysed in: DPBS (BI Biologicals) supplemented with 30mM NaCl, 10mM Hepes (pH 7.4), 1.5mM MgCl2, 0.5% Triton X-100, 5% glycerol, 10mM imidazole, 1mM PMSF and Protease Inhibitor Cocktails (Sigma: P8849). All steps were carried out at 4°C. The lysates were passed 3 times through a syringe (21G needle), clarified by centrifugation for 7 minutes at 16,000x g. The supernatants were adjusted to have equal concentration of total protein and loaded on 15μl of HisPur Ni-NTA Resin (Cat. #88221,ThermoFisher Scientific). After incubation for 40 minutes, the flow through was allowed to pass through the beads twice more, and the beads were washed five times with DPBS supplemented with 146mM NaCl, 20mM Tris-HCl (pH 8.0), 0.5% Triton X-100, 5% glycerol, 15mM imidazole. The His-tagged Bag3 along with associated proteins was eluted with 300mM imidazole in 50mM Na(PO4) (pH 6.8), 300mM NaCl. The eluted samples were immunoblotted with respective antibodies (16).

For pull-down analysis of ubiquitinated polypeptides, MCF10A (Untransformed) cells plated on 10cm dish and treated with JG-98 for 12 hours or left untreated. After that cells were washed with DPBS and fixed with 1.2% formaldehyde followed by a wash with 50mM Tris-HCl (pH 7.4) in DPBS at 4°C. The cells were lysed in: DPBS (BI Biologicals) supplemented with 30mM NaCl, 10mM Hepes (pH 7.4), 1.5mM MgCl2, 0.5% Triton X-100, 5% glycerol, 10mM imidazole, 1mM PMSF and Protease and Phosphatase Inhibitor Cocktails. All steps were performed at 4°C. The cells were lysed with the buffer: (50mM Tris-HCl (pH 7.4), 150mM NaCl, 1% Triton X-100, 5mM EDTA) supplemented with Protease Inhibitor Cocktails (Sigma: P8340) and clarified by centrifugation for 7 minutes at 16,000x g. The supernatants were adjusted to have equal concentration of total protein and then supplemented with 10μg of His6-Ubiquilin-1 Tandem UBA (BostonBiochem: UBE-110). After incubation for 60 minutes, 15μl of HisPur Cobalt Resin (Pierce, ThermoFisher Scientific) were added, and all the following steps described for the His-tagged Bag3 pull-down were performed (16).

### Data Analysis

Hermes Wiscon Imaging System with Athena Winsoft software package (V1.0.10) was used for immunofluorescence experiments. Fiji-ImageJ was used for all quantification of immunoblotting. Transcriptome analysis was done using pipeline developed inhouse, differentially expressed genes were computed through limma package (38–40). Further gene enrichment was computed using the Broad Institute software package for gene set enrichment analysis, Also, R studio packages and GraphPad Prism (v9) was used to compute statistical parameters and plotting. Statistical analysis was performed by Student’s t test, one-way analysis of variance (ANOVA), or two-way ANOVA as appropriate. Data were expressed as means ± SEM.

### RNAseq processing

RNA was extracted from cells using the RNeasy Mini kit (Cat. #74104, Qiagen). Library preparation strategy (BGISEQ-500) was adopted and performed by BGI, China.

### Data Processing

We developed our own pipeline to analyze the data where reads were first trimmed and clipped for quality control in trim_galore (v0.5.0) and checked for each sample using FastQC (v0.11.7). Data was aligned by Hisat2 (v2.1.0) using hg38, GRch38.97. High-quality reads were then imported into samtools (v1.9 using htslib 1.9) conversion to BAM file. Gene-count summaries were generated with featureCounts (v1.6.3): A numeric matrix of raw read counts was generated, with genes in rows and samples in columns, and used for differential gene expression analysis with the Bioconductor packages (38–40).

## Data availability

All data supporting the findings of this publication can be found within the supporting information and the primary publication except those noted here. The transcriptome analysis related raw FASTQ files, count file and differentially expressed gene list can be accessed via GSE171440.

## Supporting information

This article contains supporting information.

## Acknowledgement

We are grateful for the services from MD Anderson Cancer Center for Proteomics by Reverse Phase Protein Array (RPPA).

## Funding and additional information

This work was supported by NIH Grant-RA1800000163 and Israel Science Foundation Grant: ISF-1444/18, ISF-2465/18.

